# *BrphyB* is critical for rapid recovery to darkness in mature *Brassica rapa* leaves

**DOI:** 10.1101/2020.05.22.111245

**Authors:** Andrej A. Arsovski, Joseph E. Zemke, Morgan Hamm, Lauren Houston, Andrés Romanowski, Karen J. Halliday, Nathalie Nesi, Jennifer L. Nemhauser

## Abstract

Crop biomass and yield are tightly linked to how the light signaling network translates information about the environment into allocation of resources, including photosynthates. Once activated, the phytochrome (phy) class of photoreceptors signal and re-deploy carbon resources to alter growth, plant architecture, and reproductive timing. *Brassica rapa* has been used as a crop model to test for conservation of the phytochrome–carbon network. *B. rapa phyB* mutants have significantly decreased or absent CO_2_-stimulated growth responses in seedlings, and adult plants have reduced chlorophyll levels, photosynthetic rate, stomatal index, and seed yield. Here, we examine the transcriptomic response of adult wild-type and *BrphyB* leaves to darkening and recovery in light. Three days of darkness was sufficient to elicit a response in wild type leaves suggesting a shift from carbon fixation and nutrient acquisition to active redistribution of cellular resources. Upon a return to light, wild-type leaves appeared to transcriptionally return to a pre-darkness state restoring a focus on nutrient acquisition. Overall, *BrphyB* mutant plants have a similar response with key differences in genes involved in photosynthesis and light response which deviate from the wild type transcriptional dynamics. Genes targeted to the chloroplast are especially affected. Adult *BrphyB* mutant plants had fewer, larger chloroplasts, further linking phytochromes, chloroplast development, photosynthetic deficiencies and optimal resource allocation.

## INTRODUCTION

Light plays at least two distinct roles in shaping plant form and productivity. First, light is essential for photosynthesis, which allows plants to convert the energy held in photons into the high potential energy found in the chemical bonds of sugars. Second, light provides information on how a plant can optimize its architecture to maximize photosynthetic potential in a given environment. How these two light systems are coordinated remains largely unknown, especially in mature leaves.

Limited light supply by an established canopy triggers a rapid shade-avoidance response that is characterized by increased elongation growth rate of stems and petioles, decreased leaf surface area and thickness, and delayed leaf yellowing (Casal, 2012; Franklin and Whitelam, 2005). On the other hand, partial plant shading or darkening will induce a range of responses between acclimation to leaf senescence (Weaver and Amasino, 2001; Brouwer *et al.*, 2012). These processes directly reduce the impact of shade or dark while additional responses such as acclimation of the photosynthetic apparatus rather help to fine tune the use of resources under shade/dark.

Plants use an array of photoreceptors to capture and transduce the light signal in diverse responses known collectively as photomorphogenesis. Photoreceptors’ absorbance properties span most of the visible light spectrum, from the phytochromes that absorb in the red (R)/far-red (FR) to the cyptochromes and phototropins that absorb in the blue/near-ultraviolet to the UV-receptors. Among these, the phytochromes (phys) are among the best characterized (Chen *et al.*, 2004). Upon illumination, phys undergo conformational changes from an inactive (Pr) to an active (Pfr) form (Fraser *et al.*, 2016), which is subsequently translocated into the nucleus and participates in transcriptional regulation (Chen *et al.*, 2005; Castillon *et al.*, 2007). Five *PHY* genes have been described in the *A.thaliana* genome (*PHYA*-*PHYE*) with partial overlapping functions (reviewed in Chen *et al.*, 2004).

Phytochrome-dependent light signaling that initiates photomorphogenesis has been extensively studied using the seedling model (reviewed in Arsovski *et al.*, 2012). In addition, it is clear from work in *A.thaliana* that phytochromes control chloroplast gene expression, as well as nuclear-encoded factors involved in chloroplast development (Oh and Montgomery, 2014; Nevarez *et al.*, 2017). Recent studies in *A.thaliana* and *Brassica rapa* showed that adult *phyB* mutants have reduced chlorophyll levels, photosynthetic rate, and stomatal index. Work by a number of groups has connected PhyB to biomass accumulation, carbon resource management, seed yield and changes in metabolism across the plant life cycle (Yang *et al.*, 2016; Krahmer *et al.*, 2018; Arsovski *et al.*, 2018; Wies *et al.*, 2019).

To date, most of our knowledge about the roles of phytochromes in the dark-to-light transition primarily came from experiments focused on the de-etiolation process of seedlings (Li *et al.*, 2011). This has left a gap in our understanding about the role of phytochromes in light-activated transcription of genes in mature leaves. This is important because several light-regulatory mechanisms essential for photosynthetic efficiency and adaptation occur only in mature leaves. For example, Chory *et al*. demonstrated that the primary role of phytochrome in greening *A. thaliana* plants is in modulating the degree rather than the initiation of chloroplast development (Chory *et al.*, 1989).

In this study, we investigated the effects of phyB on gene expression upon dark-to-light transition in the mature leaf of *B. rapa* by comparing the transcriptomic responses between wild-type and a *phyB* mutant. *B. rapa* is closely related to *A.thaliana* (Wang *et al.*, 2011) but its leaves are significantly larger. Larger leaves cause more self-shading, and, in combination with the longer life of *B. rapa* compared to *A.thaliana*, there is more total demand for resources. As the *B. rapa* genome contains only one *PhyB* ortholog and no likely ortholog for the closely related *AtPhyD*, we took advantage of the *BrphyB3* mutant allele described previously (Arsovski *et al.*, 2018). Wild-type and *BrphyB* leaves exhibited significant overlaps in their transcriptomic response to dark and recovery; however, gene ontology analyses pointed out important misregulations in *BrphyB* mutant for genes involved in nitrogen metabolism, light harvesting and photosynthesis. Altogether these results support a role for PhyB in chloroplast development and resource allocation, and have implications for increasing the resource-efficiency of Brassica crops.

## MATERIALS AND METHODS

### Growth conditions of *B. rapa* adult plants

The *B. rapa* wild-type R-o-18 and *BrphyB* mutants were originally from the John Innes Center’s RevGenUK resource. The BrphyB-3 previously described in Arsovski *et al.*, 2018 was used for RNAseq experiments. BrphyB-1 was also previously described in Arsovski *et al.*, 2018. Seeds were planted directly into our standard soil mix of 1:1 Sunshine Mix #4 (SunGro Horticulture):vermiculite. Plants were grown in 2.6 liter square pots (McConkey Grower Products; Sumner, WA, USA) and bottom-watered daily in long day conditions (16 h light, 8 h dark, ~115 μmol.m^−2^.s^−1^ light intensity) in a Percival E-30B growth chamber (https://www.percival-scientific.com/) set to 20°C. Experiments were conducted at 3 weeks and the plants were then moved to growth room until seed harvest.

### Leaf sample preparation

Three weeks after sowing, two developmentally matched leaves from each wild-type and *BrphyB-3* plant were tagged that corresponded to the first and second true leaves. Samples were collected at ZT 5 using a standard hole punch (28 mm^2^ circular area of leaf blade tissue) with symmetrical harvest (a second hole punch on the other side of the other side of the mid-vein of the same leaf) for chlorophyll assay, chloroplast measurements and transcriptome analysis. Tissue from 3 individual plants was combined to make one biological replicate. At 3 weeks of age the “Pre” sample was collected from the first leaf while the second one was covered with tinfoil. 24 hours later the “24hr” sample was harvested from the uncovered leaf, the same leaf that provided the “Pre” sample. Then, 48 hours later the tinfoil was removed from the covered leaf and the “dark” samples were similarly collected. Finally, 24 hours later the “recovery” samples were collected from this same leaf. Samples were immediately flash frozen in liquid nitrogen (Fig.1). In total, three biological replicates were collected in similar fashion.

**Figure 1:**
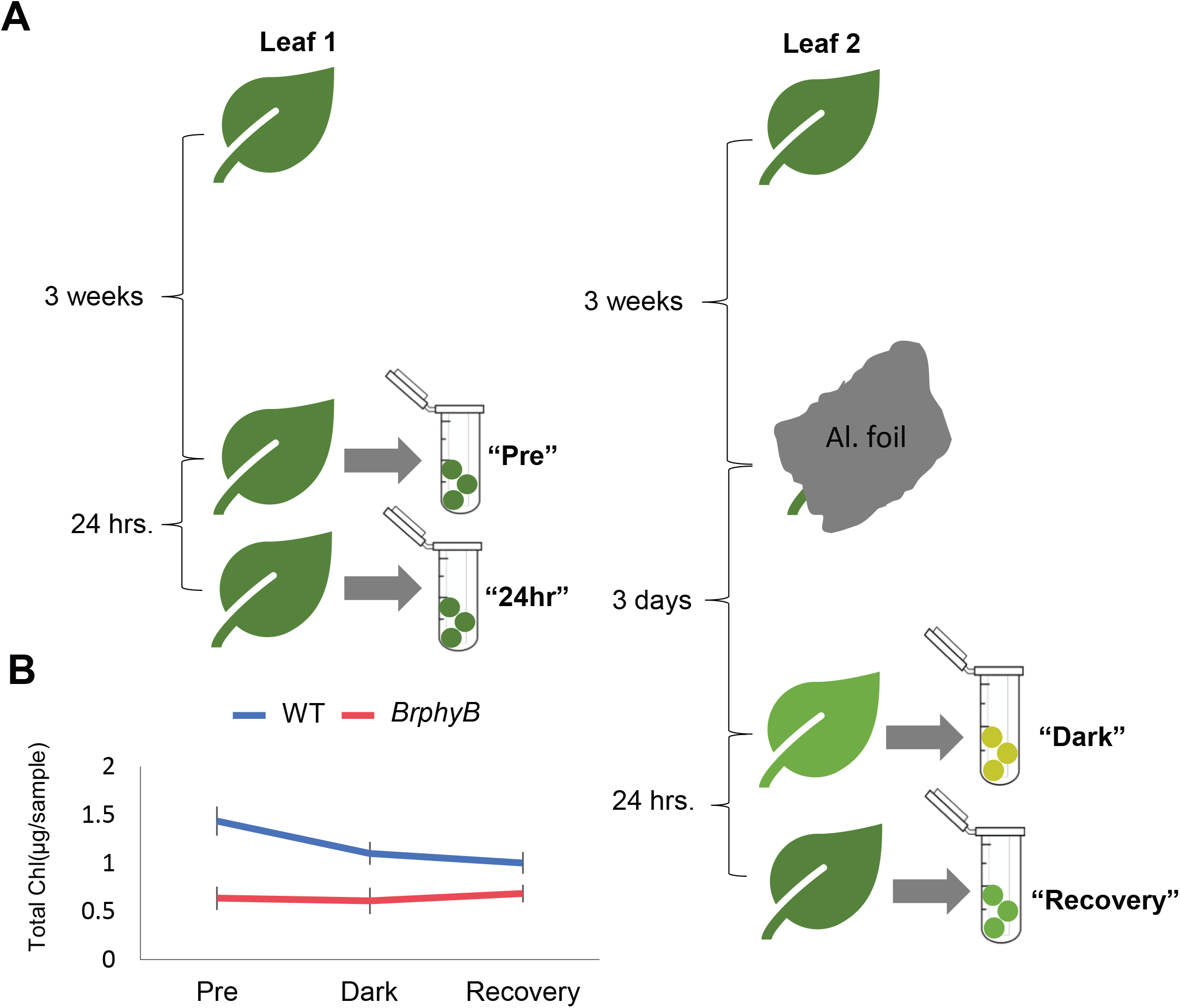
RNAseq experimental set-up (A). At 3 weeks of age the “pre” sample was collected from the first of the two developmentally matched leaves. Symmetrical samples from the same leaf was collected for chlorophyll measurement. The second matched leaf was covered with tinfoil at this time. 24 hours later the “24” sample was harvested from the uncovered leaf, the same leaf that provided the “pre” sample. 48 hours later the tinfoil was removed from the covered leaf and the “dark” samples were similarly collected. 24 hours later the “recovery” samples were collected from this same leaf. Samples were immediately frozen in liquid nitrogen. Three biological replicated were similarly collected. Total chlorophyll in Pre, dark, and recovery samples, error bars are SE.

### Chlorophyll measurement

For chlorophyll measurement, ethanol extractions were done as in (Yang *et al.*, 2016). Determinations were run by measuring optical density at 645 nm and 665 nm using an Epoch Microplate Spectrophotometer (www.biotek.com). Values were obtained using the following formulas: Chl *a*=5.21*A*_665_−2.07*A*_645_; Chl *b*=9.29*A*_645_−2.74*A*_665_, for Chlorophyll A and B, respectively. Three individual biological replicates were used for this assay.

### RNAseq

Leaf tissue was disrupted with Zirconia/Silica beads for 1 minute in a MiniBeadbeater-96 (BioSpec Products, Inc.) while frozen. After adding 500 μL of Lysis/Binding Buffer to each sample and vortexing until homogeneous, samples were run on the MiniBeadbeater-96 for an additional minute. Following tissue disruption samples were centrifuged at 16,000 × g for 10 minutes at 20 °C. For each sample, a 50 μL aliquot of the supernatant was added to 50 μL of NEB RNA binding buffer and mRNA isolated as per the NEBNext® Poly(A) mRNA Magnetic Isolation Module manual.

RNA-seq libraries were prepared by using the Full Transcript mode YourSeq Dual (FT & 3’-DGE) RNAseq Library Kit (Amaryllis Nucleics). A Bioanalyzer 2100 (Agilent, High Sensitivity DNA Kit) was used for library quality control, to determine average library size, and together with concentration data from a Qubit 2.0 Fluorometer (Life Technologies, dsDNA High Sensitivity Assay Kit) to determine individual library molarity and pooled library molarity. Pooled libraries were sequenced on a NextSeq 500 (Illumina, High Output v2 75 cycle kit) to yield single-read 80 bp reads.

FASTQ processing was performed by Amaryllis Nucleics (Oakland, CA). Sequence files were preprocessed in two steps. A Python library (clipper.py, https://github.com/mfcovington/clipper) was used to trim off the first 8 nucleotides of each read to remove potential mismatches to the reference sequence caused by annealing of a random hexamer required for library synthesis. Trimmomatic v0.36 (http://www.usadellab.org/cms/?page=trimmomatic) was used to remove adapter sequences and trim or filter reads based on quality The parameters used for Trimmomatic were ILLUMINACLIP:TruSeq3-PE-2.fa:2:30:10 LEADING:3 TRAILING:3 SLIDINGWINDOW:4:15 MINLEN:50.

Preprocessed reads were mapped to the *Brassica rapa* v2.5 genomic reference sequence (http://brassicadb.org/brad/datasets/pub/Genomes/Brassica_rapa/V2.0/V2.5/Chr/BrapaV2.5_Chr.fa.gz) using bowtie2. Read counts for each gene in the gene annotation file (http://brassicadb.org/brad/datasets/pub/Genomes/Brassica_rapa/V2.0/V2.5/Chr/BrapaV2.5_Chr.gene.gff.gz) were calculated using htseq-count (with the −s yes parameter to enforce strand-specific alignment) from the HTSeq Python library (https://academic.oup.com/bioinformatics/article/31/2/166/2366196; http://htseq.readthedocs.io/en/master/index.html).

The package edgeR (Robinson *et al.*, 2010) was used to process the expression matrix and identify differentially expressed genes between treatments and genotypes. For the main analysis, the generalized linear model functionality of this package, based on a negative binomial distribution model for gene expression was used to identify differentially expressed genes. Genes were considered significantly differentially expressed based on having a fold change greater than 2-fold up or down between conditions, and a q-value (adjusted p-value by Benjamini-Hochberg procedure - (Benjamini and Hochberg, 1995) less than 0.01.

Contrasts between the Pre and 24hr timepoints were used to identify genes that could be exhibiting differential expression caused by wound response from the tissue sampling rather than response to darkness and recovery. Genes identified in these wound control contrasts were tagged but not excluded from the rest of the analysis. Dispersion was estimated independently for the wound control contrasts based on the Pre and 24hr timepoints only. This was done because using the dispersion estimates from the main analysis, including the Dark and Recovery timepoints resulted in biased p-value distributions due to the significant change in expression of most genes in the dark timepoint as compared to the other timepoints (68% of genes between wild type-Dark and wild type-Pre).

Complete data can be accessed from the Gene Expression Omnibus (GEO) under the entry GSE135955.

### Venn diagrams and gene ontology (GO) analysis to prioritize the DEG

To find differences between the wild-type and *phyB* response to darkness and reillumination we looked at sets of genes that were significantly changed in these contrasts (darkened vs. pre or recovery vs. darkened) in one genotype but not the other. For each up or down Venn-diagram used in the GO analysis, all genes which either the wild-type or *BrphyB* mutant were greater (or less for down regulated Venn) than the log fold change cutoff of 2 were considered. These genes were then put into 4 categories: wild-type significant only, *BrphyB* significant only, both wild-type and *BrphyB* significant, and neither genotype significant.

To obtain a comprehensive list of all *B. rapa* presumptive orthologs in *A. thaliana* we used each *B. rapa* protein as a query to perform local homology searches. Briefly, for each protein in the *B. rapa* v2.5 proteome, the best *A. thaliana* hit was retrieved by sequence similarity search using a local installation of the BlastP algorithm (Protein-Protein BLAST 2.7.1+) of the NCBI tool BLAST (http://blast.ncbi.nlm.nih.gov/Blast.cgi) against the A. thaliana ARAPORT11 proteome, with default parameters and the outfmt parameter set to 7 (to obtain a tabular output with comment lines). This resulted in a collection of the best A. thaliana hits for each *B. rapa* protein. This output was further processed to limit each ortholog to the best hit, using a custom BASH script (available upon request). GO term enrichment analysis was performed against the well annotated *A.thaliana* genome using the PANTHER classification System (v.14.1 available at http://pantherdb.org/; (Mi *et al.*, 2019).

### Law 2018 comparison

For the data in Table S3, we created a list by matching the *A. thaliana* gene IDs from the (Law *et al.*, 2018) analysis up to the *B. rapa* genes they were found to be the nearest homolog of (see GO analysis section above). We considered the genes that were differentially expressed after 3 days (D3) of darkness applied on an individualized leaf (IDL) or a whole plant (DP). *A. thaliana* genes that did not match to *B.rapa* genes in our set were dropped from the comparison, *A. thaliana* genes with multiple brassica homologs were listed multiple times in this list. The elements of this list were then broken down into a Venn diagram based on whether they were considered significantly differentially expressed up or down in our darkened vs. pre contrast and Law’s IDL_D3 and DP_D3 contrasts. Each category list was then reduced to only contain unique *A. thaliana* IDs. These counts are displayed in Table S3.

### RNA extraction and quantitative real-time PCR (qRT-PCR) analysis

Expression analysis was performed using 4 biological replicate samples collected identically as described for RNA sequencing. Each sample was immediately frozen in liquid nitrogen and stored at −80 °C until processing. Frozen tissue was ground in liquid nitrogen and total RNA was extracted using the GE Illustra RNA kit (GE Life Sciences), and 2 μg of eluted RNA was used for cDNA synthesis employing iScript (Bio-Rad). Samples were analyzed using SYBR Green Supermix (Bio-Rad) reactions run in a C100 Thermal Cycler (Bio-Rad) fitted with a CFX96 Real-Time Detection System (Bio-Rad). Relative expression levels were calculated using the formula (E_target_)^−CPtarget^/(E_ref_)−^CPref^ (Pfaffl, 2001) and normalized to the *B.rapa PP2A* (Brara.F00691) reference gene. qPCR primer sequences are as follows: *BrPP2A* (forward 5’-TCGGTGGTAACGCCCCCGAT-3’; reverse 5’-CGACTCTCGTGGTTCCCTCGC-3’); *BrMGT6* (Brara.E02300) (forward 5’-CAGCATCCGCCACCGCAAGA-3’; reverse 5’-GCCTTCGCAACAACCGCAGC-3’); *BrRLK*4 (Brara.I00004) (forward 5’- TCCGCCGTCGCGATCTCTCT-3’; reverse 5’-CCCGCTCCAAACGCTTGTCCA-3’); *BrPIFI* (forward 5’-GCCACCACTCTTGCACCCCC-3’; reverse 5’- CCGCGGTTGGAGGAAGACCG-3’).

### Code

The R code used to generate all the analysis results is provided in supplement X and can also be found on github.com/nemhauser-lab/brassica_rna_seq.

### Methods, Motif Enrichment

A set of binding motifs for 619 *A.thaliana* transcription factors was downloaded from plantTFDB (http://planttfdb.cbi.pku.edu.cn/), (Jin *et al.*, 2014, 2015, 2017). The RSAT matrix clustering tool (http://rsat.eead.csic.es/plants/matrix-clustering_form.cgi (Castro-Mondragon *et al.*, 2017) was used with default parameters to group the motifs into 56 clusters based on similarity of aligned position weight matrices. Genes that were significantly differentially expressed from conditions pre to dark, and from dark to recovery were divided into groups depending on 3 factors: genotype, direction of regulation (up or down), and whether they are annotated with the GO term “chloroplast” (GO:0009507). The package “motifmatchr” (Schep, 2019) was used to count the number of genes in each set with promoter sequences with matches to each of the 619 motifs. Promoters sequences were defined as the 1000 base pairs immediately upstream of the gene start position as defined by the genome annotation file. The fisher exact test was then used to determine if there was significant enrichment for each motif between genes only significanty regulated in WT and genes only significantly regulated in *BrphyB* in these sets. The raw p-values from these tests were adjusted using the by Benjamini-Hochberg procedure (Benjamini and Hochberg, 1995). Adjusted p-values of <0.01 were considered significant.

All motifs found significant from these tests closely matched to three canonical motifs described in the literature: the G-box motif CACGTG, the Evening Element AGATATTTT, and the Telo-box motif AAACCCTAA. The proportions of promoters with one or more exact matches to three canonical motifs were found for the gene sets described above.

### Chloroplast Measurement

Tissue was immediately cleared after collection and fixed using ClearSee solution as described in (Kurihara *et al.*, 2015). Images were taken using a Leica TCS SP5 II laser scanning confocal microscope (https://www.leica-microsystems.com). Chloroplast number, area, and density were determined using ImageJ software.

## RESULTS

### Mature *BrPhyB* mutant leaves have significant transcript reductions of chloroplast targeted genes

Loss of phyB leads to significant reductions in both chlorophyll levels and rates of photosynthesis in three-week-old *B. rapa* plants (Arsovski *et al.*, 2018). To further understand the link between phyB, chloroplast development, and photosynthesis we examined the transcriptomic response of mature leaves that were subjected to three days of darkness before being reintroduced into the light. As part of this experiment we first compared the transcriptome of three-week-old *B.rapa* wild type and Br*phyB* leaves. Genes were considered differentially expressed if the fold change between timepoints or genotypes was greater than 2, and the significance (adjusted p-value) was less than 0.01. 114 genes were significantly upregulated in *BrphyB* leaves compared to wild type. Unsurprisingly, these include *B.rapa* orthologs to *A.thaliana LONG HYPOCOTYL IN FAR-RED*(*HFR1*), *PHYTOCHROME INTERACTING FACTOR 3-LIKE 1 (PIL1), PHYTOCHROME-INTERACTING FACTOR 6 (PIF6)*, and *INDOLE-3-ACETIC ACID INDUCIBLE 29(IAA29).* 79 genes were significantly downregulated in Br*phyB* leaves compared to the wild type. Gene Ontology (GO) analysis of cellular location annotations showed a strong enrichment for the chloroplast envelope, stroma, and photosystem II. These include *B.rapa* orthologs for *A.thaliana PHOTOSYSTEM II SUBUNIT P*, *PHOTOSYSTEM II BY*, *LIGHT-HARVESTING CHLOROPHYLL B-BINDING PROTEIN 3*, and *CHLOROPHYLL A/B BINDING PROTEIN 1* (Table S1).

### Darkening of individual leaves for three days initiates resource reallocation

Samples were taken from the first or second true leaf of three-week-old wild-type and *BrphyB*-3 plants (hereafter termed “pre”). As a wounding control, a second sample was taken from the “pre” leaves 24 hours later (hereafter termed “24hr”). Leaves that were developmentally-matched with those selected for the “pre” treatment were covered with foil. After three days, the foil was removed and the “darkened” sample was collected immediately. The “recovery” sample was collected from the same leaf 24 hours after this timepoint to capture the earliest stages of recovery (Fig. 1A). Matching samples were collected from each leaf to assay chlorophyll levels. At three weeks old, *BrphyB* mutants are visibly paler compared to same aged wild-type plants, and have significantly reduced chlorophyll levels (Fig. 1B and Arsovski *et al.*, 2018). Three days of dark resulted in a 23% reduction of chlorophyll levels in wild type leaves while levels remained low in the mutant. This is consistent with similar experiments performed on individually darkened *A.thaliana* leaves where total chlorophyll levels and protein decline was observed after two days of darkness (Weaver and Amasino, 2001). The 24 hours of light exposure for the recovery samples was not sufficient to restore chlorophyll levels in either wild-type or *BrphyB*-3 leaves (Fig. 1B).

Extended darkness of leaves acts as a signal to initiate the organized breakdown and remobilization of valuable resources to growing vegetative and reproductive tissues (Himelblau and Amasino, 2001; Buchanan-Wollaston *et al.*, 2003; Lim *et al.*, 2007). We performed RNAseq analysis on the pre, 24hr, dark and recovery samples to assess the specific response to darkness and return to light of mature leaves in *B. rapa*. We began our analysis with the response to darkness, as previous studies in *thaliana* had already shown that dark stress is accompanied by dramatic transcriptional changes, as well as depletion of chlorophyll and large-scale degradation of proteins (Guo *et al.*, 2004; Keech *et al.*, 2007; Law *et al.*, 2018). The expression of 6852 *B.rapa* genes was significantly altered in leaves after three days in darkness. Gene Ontology (GO) analysis of predicted *A. thaliana* orthologs showed a pattern consistent with the overall expectations of metabolic reprogramming seen in other species. The 3110 genes up-regulated in response to dark were mainly involved with autophagy, catabolism, leaf senescence and vesicle fusion (Fig. 2A; Table S2). Conversely, down-regulated genes were mainly involved in photosynthesis, biosynthetic processes and plastid translation. Together, these data suggest a shift from carbon fixation and nutrient acquisition to active redistribution of cellular resources (Fig. 2B, Table S2).

**Figure 2:**
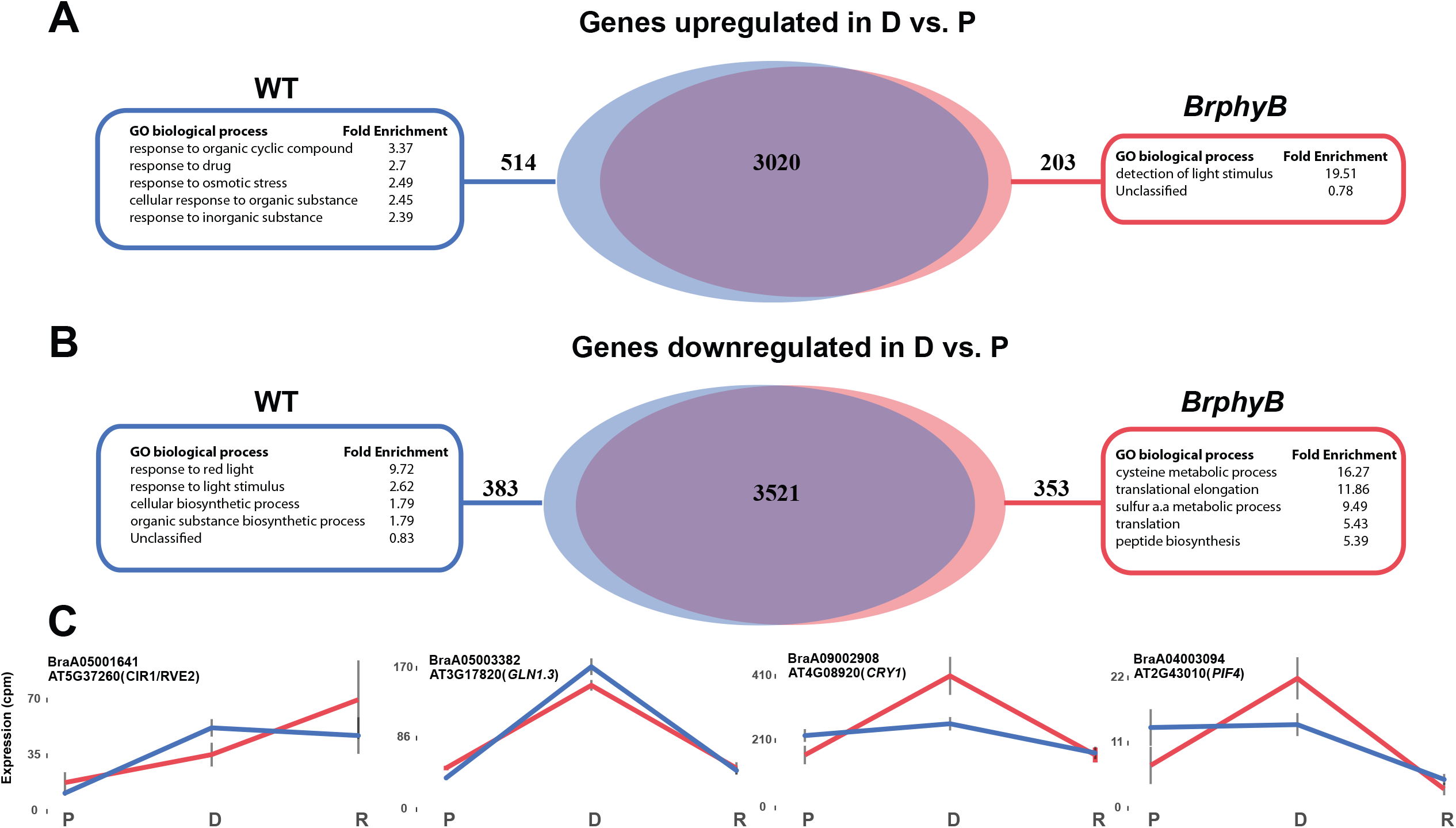
Genes differentially expressed in dark. A Gene Ontology (GO) analysis of genes uniquelly differentially expressed in wild type and *BrphyB* mutant leaves following 72 hours of dark. A) Upregulated genes. B) Downregulated genes. C) Expression values of 3 biological replicates in exemplar genes in Pre(P), Dark (D) and Recovery (R).

A recent experiment in *A.thaliana* found that the effect of darkening individual leaves was substantially similar to the effect of darkening whole plants, albeit with distinct timing for peak differences in gene expression between the two treatments Law *et al.,* 2018). When we compared our transcriptomic response to this dataset, we found a substantial overlap. The highest similarity between the *B.rapa* dark response was to *A.thaliana* individually darkened leaves for 3 days (IDL_3D). Of the genes significantly up or down regulated in *A. thaliana* individually darkened leaves after 3 days, 46.7% had *B.rapa* homologs also significantly up or down regulated (in the same direction) (Table S3). Shared *A. thaliana* genes upregulated in response to dark were significantly enriched for GO terms such as autophagy, catabolic process, and leaf senescence (Table S4). The darkening response in *B.rapa* was more similar to that of individually darkened leaves than whole darkened plants in *A.thaliana*. Of the genes found to be up/down regulated in individual leaves (IDL_D3) but not whole darkened plants (DP_D3), 35.3% had *B.rapa* homologs with significant change in the same direction, compared to 25.1% in whole darkened plant unique genes (Table S3,4). In *A. thaliana*, 167 senescence-associated genes were shown to change expression in response to darkening (Parlitz *et al.*, 2011). The *B.rapa* orthologs of 103 of these genes do not show significant changes in response to dark. Of the remaining 57 senescence-associated genes, 42 show a reversible pattern of upregulation in the dark and downregulation upon re-illumination. 15 are ‘non-reversible’, upregulated in the dark without significant changes upon a return to light (Table S5).

Many of the genes that were regulated by returning the leaves to light were similar to those already identified as light-responsive from experiments in seedlings. In *A. thaliana*, expression of up to one-fourth of the whole genome is altered in seedlings grown for 4 days in red light compared to those grown in the dark (Shi *et al.*, 2018). These changes are largely mediated by a small group of transcription factor families which include the PHYTOCHROME INTERACTING FACTORS (PIFs). Nearly 60% of PIF-dependent, red light induced genes in *A. thaliana* seedlings have Gene Ontology (GO) annotations indicating functions related to photosynthesis and chloroplast (Leivar *et al.*, 2012). In *B.rapa* wild type leaves 3756 genes were upregulated in the recovery condition when compared to the dark timepoint. The most significantly enriched GO terms were response to light stimulus, photosynthesis, translation and metabolism, suggesting a return to a pre-darkness transcriptional state (Figure 3A, Table S2). The 3299 genes downregulated in recovery compared to dark were mainly involved in catabolism, vesicle fusion and transport, and protein degradation further supporting a shift from resource remobilization towards nutrient acquisition (Figure 3B, Table S2).

**Figure 3:**
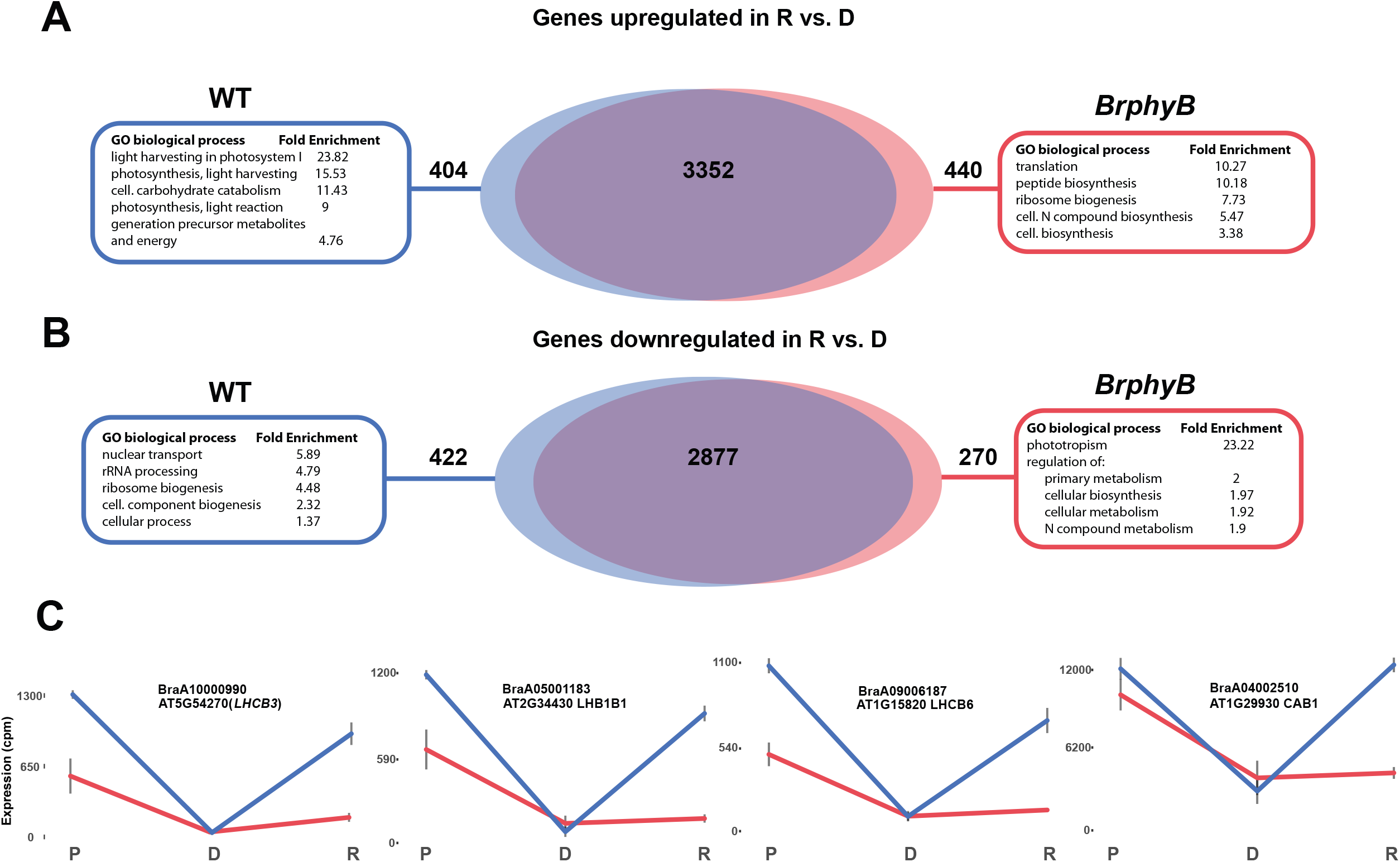
Genes differentially expressed on return to light. A Gene Ontology (GO) analysis of genes uniquelly differentially expressed in wild type and BrphyB mutant leaves 24 hours after return to light. A) Upregulated genes. B) Downregulated genes. C) Expression values of 3 biological replicates in exemplar genes in Pre(P), Dark (D) amd Recovery (R).

### *BrphyB* is critical for full recovery response

RNAseq analysis of *BrphyB* individually darkened leaves revealed an essentially similar response to what was observed in wild-type plants. 7,994 genes were differentially expressed in wild-type and *BrphyB*-3 leaves after 3 days of dark treatment compared to the pre samples. The vast majority (81.8%) of this response was shared between the genotypes (Fig. 2A, B). However, analysis of GO terms significantly enriched in the uniquely wild-type or *BrphyB* differentially regulated gene sets illustrated phyB-dependent responses to darkness. Unique wild-type enriched terms were largely related to cellular responses to organic and inorganic compounds, drugs and stress (Table S6). A closer look at these phyB-controlled groups revealed orthologs to *A. thaliana* genes REVEILLE2, REVEILLE8, and CIRCADIAN CLOCK ASSOCIATED1—all key transcriptional regulators of circadian rhythm, auxin and stress response in *A. thaliana* and known to act downstream of PhyB (Fig. 2A,C) (Alabadí *et al.*, 2001; Zhang *et al.*, 2007; Rawat *et al.*, 2011; Farinas and Mas, 2011; Jiang *et al.*, 2016). *BrphyB* unique up-regulated genes were enriched in genes associated with response to light, including *B.rapa* orthologs of PHYTOCHROME INTERACTING FACTOR4, PHYTOCHROME INTERACTING FACTOR5, PHYTOCHROME KINASE SUBSTRATE1 and CRYPTOCHROME 1 (Table S6, Fig. 2C).

PhyB-repressed genes are enriched for categories such as response to light stimulus and cellular biosynthetic process, while those activated by phyB are enriched for categories related to protein synthesis such as cysteine metabolic process, translational elongation, and peptide biosynthesis (Fig.2B). These include *rapa* orthologs to three *A. thaliana* glutamate-ammonia ligases (GLUTAMINE SYNTHASE 1;2, 1;3 and 1;4) with roles in nitrogen remobilization and seed yield (Guan *et al.*, 2015), stress response and pollen viability (Ji *et al.*, 2019). In *B. rapa*, *BrphyB* mutants have up to a 90% decrease in seed yield (Arsovski *et al.*, 2018); however, we did not observe a difference in area or weight of seeds compared to wild type (data not shown). This would suggest that plants are re-calibrating the amount of resources available, and maintaining quality by partitioning them into a smaller number of seeds.

While there is substantial overlap between the response of wild type and *BrphyB* mutants to re-illumination (80.2% of the 7,765 genes are common to both genotypes), there are also several key differences. To validate the RNAseq results, we selected three genes whose expression in wild type was significantly downregulated during darkening compared to pre followed by a significant upregulation in recovery compared to darkening. qPCR of these genes was done in wild-type, *BrphyB*-3, and an additional mutant allele *BrphyB*-1. Brara.E02555 is an ortholog of the *A.thaliana* At3g15840 gene. In *A.thaliana* POST-ILLUMINATION CHLOROPHYLL FLUORESCENCE INCREASE (PIFI) is a nuclear-encoded chloroplast protein essential for NDH-mediated non-photochemical reduction of the plastoquinone pool in chlororespiratory electron transport (Wang and Portis, 2007). The *A.thaliana* orthologs RECEPTOR-LIKE PROTEIN KINASE 4 (RLK4) and MAGNESIUM TRANSPORTER 6 (MGT6) are a Ser/Thr receptor-like protein kinase expressed in the root and a magnesium transporter required for growth in Magnesium limited conditions, respectively. The qPCR expression closely resembled the RNAseq results for wild-type and *BrphyB*-3 and *BrphyB*-1 expression matched that of *BrphyB*-3 for all three genes. *BrPIFI* expression decreases dramatically in response to darkening in both wild-type and *BrphyB*-1 mutant leaves. However, while *BrPIFI* expression increases in response to the leaf’s return to light in wild-type, it remains low in the mutant. Similarly, *BrRLK4* and *BrMGT6* expression increases in recovery in wild-type leaves. However, in *BrphyB*-3 leaves expression of both decreases 24 hours after the cover is removed from the leaf (Fig. S1) (Coello *et al.*, 1999; Wang and Portis, 2007).

BrphyB is required for rapid return to the full photosynthetically-active transcriptional program. There are 404 genes up-regulated in recovery of only wild-type leaves. These genes are mainly involved in light harvesting in Photosystem I, photosynthesis, cellular carbohydrate catabolism and generation of precursor metabolites and energy and are not upregulated in *BrphyB* leaves 24 hours after return to light (Fig. 3A, Table S6). We previously showed that the expression *B.rapa* GOLDEN2-LIKE 1 (BrGLK1) increases 70% in response to high CO^2^ in wild type seedlings but decreases in *BrphyB* mutants (Arsovski *et al.*, 2018). In *A.thaliana* GLK1 is one of a pair of partially redundant transcription factors that affect the expression of nuclear photosynthetic genes involved in chloroplast development (Waters *et al.*, 2008, 2009; Kobayashi *et al.*, 2013). Here the *B.rapa* ortholog of *A.thaliana* GOLDEN2-LIKE 2 is significantly upregulated upon return to light in wild type but not *BrphyB* mutant leaves (Table S7). Closer examination of chloroplast localized genes whose expression significantly changes in response to darkening and recovery revealed a stark contrast in responsiveness between the two genotypes. In response to darkening, 1861 chloroplast related genes were significantly downregulated in either genotype. Of these, 94.7 % (1763 genes) were unique to wild type leaves and were not significantly downregulated in the mutant. Upon a return to light 1904 chloroplast localized genes were significantly upregulated with 85.8% common to both genotypes. However, 131 localized genes were upregulated only in wild type leaves while 140 were unique to the *BrphyB* mutant. GO cellular component analysis identified 125 genes with predicted chloroplast-localization that are up-regulated in wild-type but not BrphyB leaves during recovery. *A.thaliana* Photosystem II genes such as LIGHT-HARVESTING CHLOROPHYLL B-BINDING PROTEIN 3 (BraA10000990), LIGHT HARVESTING COMPLEX PHOTOSYSTEM II SUBUNIT 6 (BraA09006187), LIGHT-HARVESTING CHLOROPHYLL-PROTEIN COMPLEX II SUBUNIT B1(BraA05001183) and three orthologs to *A.thaliana* CHLOROPHYLL A/B BINDING PROTEIN 1 (BraA04002510, BraA07001020, BraA08002753) are upregulated on return to light only in wild type leaves (Figure 3C, Table S7).

Downregulated genes unique to wild type are enriched in annotations associated with nuclear transport, ribosome biogenesis, and rRNA processing, while terms including phototropism and regulation of primary metabolism, cellular biosynthesis and nitrogen compound metabolism are enriched in the *BrphyB* unique downregulated genes (Fig. 3B). This overall pattern suggests that *BrphyB* may be required for effective monitoring and switching between carbon- and nitrogen-demanding processes, and that this role may be essential for maximal reallocation of resources to developing seeds.

### *BrphyB* leaves have regulatory motif differences in chloroplast related genes and fewer, larger chloroplasts

In the pre condition, the 79 genes significantly downregulated in *BrphyB* leaves compared to the wild-type were enriched for localization to the chloroplast envelope, stroma, and photosystem II (Table S1). The recovery condition created a sensitized environment to detect the more immediate impacts of *BrphyB* on establishing or maintaining the photosynthetic machinery. To investigate whether there were regulatory differences in recovery between chloroplast and non-chloroplast genes in wild-type and *BrphyB* leaves we examined the promoters (1Kb upstream for TSS) of up and downregulated genes in recovery compared to dark. Genes with the GO term 0009507: chloroplast were designated as ‘chloroplast’ and those without it ‘non-chloroplast’.

The frequency of three major motifs appeared to change in response to dark and in recovery and between genotypes (Fig.S2A). The G-box element (CACGTC) is a focal point of light-regulated gene expression. In vitro gel-shift, random DNA-binding selection, and chromatin immunoprecipitation (ChIP) assays in *A.thaliana* show that four PIFs (PIF1, PIF3, PIF4, and PIF5) bound to either a G-box (CACGTG) and/or an E box (CANNTG) (Martínez-García *et al.*, 2000; Huq and Quail, 2002; Huq *et al.*, 2004; Hornitschek *et al.*, 2012). PIFs can also interact with other transcription factors at the G-box, and these interactions modulate the PIF DNA-binding activity. PIF3 and PIF4 interact with BRASSINAZOLE-RESISTANT 1 (BZR1) and bind to the same G-box DNA sequence element to regulate genes involved in the light and brassinosteroid pathways (Oh *et al.*, 2012; Zhang *et al.*, 2013). PIF1 and PIF3 also interact with the light-regulated activator ELONGATED HYPOCOTYL (HY5) at the G-box where it can both promote PIF1/3 binding and compete for binding sites (Chen *et al.*, 2013; Toledo-Ortiz *et al.*, 2014). When *B.rapa* leaves were returned to light, 37% of chloroplast genes significantly upregulated in wild type but not *BrphyB* leaves have a G-box in their promoter region compared to only 15% of non-chloroplast genes. This is not the case with chloroplast genes upregulated only in *BrphyB*. For these genes, there was essentially no difference in the number of genes a G-box whether or not they were annotated as chloroplast-associated (chloroplast genes: 13%, non-chloroplast genes: 16%) (Fig.S2B).

Differences were also present in the frequencies of Evening Element (AAAATATCT) and Telobox motif (AAACCCTAA) in chloroplast-annotated genes between wild-type and *BrphyB* leaves in recovery as well. The Evening Element (EE) motif is central to circadian clock function and environmental and endogenous signal coordination in *A.thaliana*. Key regulators of the circadian clock CIRCADIAN CLOCK ASSOCIATED1 (CCA1), LATE ELONGATED HYPOCOTYL (LHY) and REVEILLE 8 (RVE8) bind and regulate genes with EEs in their promoters (Harmer and Kay, 2005; Hsu *et al.*, 2013). Among chloroplast-annotated genes, the EE motif was present in promoter regions of 37% of genes which were significantly downregulated in *BrphyB* but not wild type leaves, while only 7% of those downregulated in wild type but not *BrphyB* had the same motif (Fig.S2C).

Short interstitial telomere motifs (telo boxes) are short sequences identical to plant telomere repeat units. In *A.thaliana* and *O.sativa* genomes telo boxes are associated with genes involved in the biogenesis of the translational apparatus (Gaspin *et al.*, 2010). Telo box motifs were enriched in the promoters of genes significantly downregulated in wild type but not *BrphyB*, and genes significantly upregulated in *BrphyB* but not wild type in recovery,15% compared to 20%, respectively. Whereas, of genes that were upregulated in wild type only, and genes downregulated in *BrphyB* only in recovery, 5% and 7% respectively had Telo-box motifs in their promoters. The differences between genes with and without the chloroplast GO annotation was less noticeable for this motif than the other two. Together these results point to a significant difference in the cis-regulatory landscape of *BrphyB* leaves (Fig. S2D).

The chloroplast carries out many functions beyond photosynthetic carbon fixation that are essential for metabolic homeostasis, including fatty acid synthesis and fixation of nitrogen and sulfur (Lopez-Juez and Pyke, 2005). Mutants with reduced phy function have significantly lower chlorophyll levels in *A.thaliana* and *B.rapa* (Ghassemian *et al.*, 2006; Strasser *et al.*, 2010; Hu *et al.*, 2013; Arsovski *et al.*, 2018). It has been suggested that phyA is primarily responsible for chloroplast maturation during de etiolation in *A.thaliana*, although there are some reports that phyB might also be involved (McCormac and Terry, 2002; Xu *et al.*, 2019).

Our results, in combination with our earlier findings that *BrphyB* mutants had reduced chlorophyll levels and photosynthetic rates, led us to hypothesize that BrphyB might be required for normal chloroplast development. We found that chloroplast density was significantly decreased in the mature leaves of the *BrphyB* mutants. Wild-type leaves had an average of 466 chloroplasts per 0.5mm^2^ compared to 326 and 253 in *BrphyB*-1 and *BrphyB*-3, respectively (Fig. 4A, B ANOVA and Tukey HSD multiple comparison test). Chloroplast area however was significantly larger in *BrphyB*-3 and *BrphyB*-1 compared to wild type, 33.6 and 41.4 to 31.3 um^2^ respectively (Fig. 4C ANOVA and Tukey HSD multiple comparison test). In *A.thaliana*, an investigation into photosynthetic, biochemical, and anatomical traits of accumulation and replication of chloroplasts (arc) mutants found that fewer, enlarged chloroplasts were less efficient at photosynthesis than more, smaller chloroplasts. Photosynthetic rate and photosynthetic nitrogen use efficiency were significantly lower in the mutants than their wild-types likely due to decreases in mesophyll conductance and chloroplast CO_2_ concentration (Xiong *et al.*, 2017). These functional differences could explain the reduced ability of *BrphyB* leaves to rapidly switch metabolic functions when exposed to darkness and again with the return to light.

**Figure 4 :**
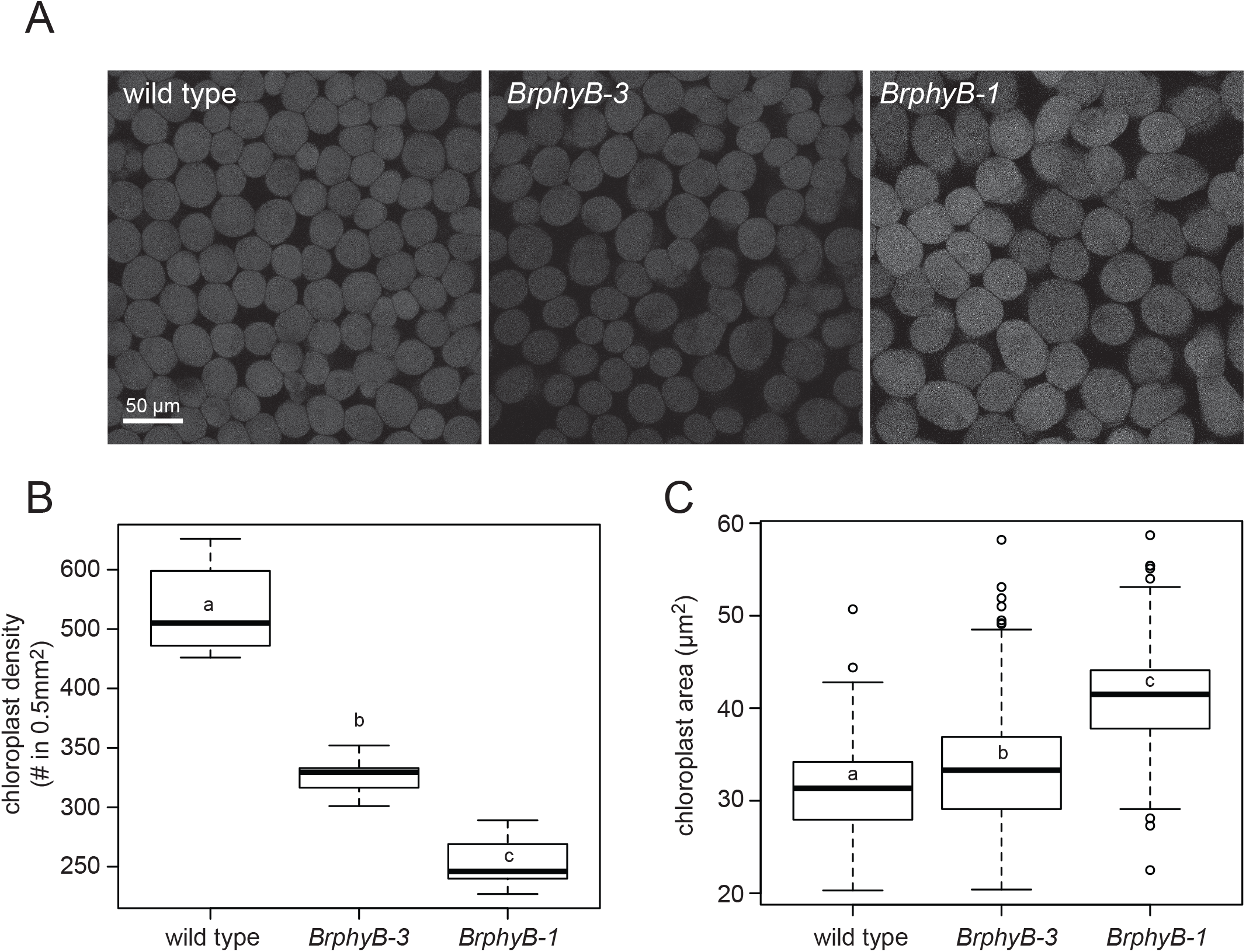
BrphyB mutant plants have fewer and larger chloroplasts. A. Fluorescent images of chloroplasts in 3 week old *B.rapa* leaves. B. Chloroplast density in same leaves as A. Chloroplast area of individual chloroplasts in same plants as A. Lower case letters in B and C indicate significant difference (ANOVA and Tukey HSD multiple comparison test; p<0.001)

## DISCUSSION

Human-driven climate change, and the associated changes in temperature, atmospheric CO_2_, and precipitation, are an urgent challenge for plant life on Earth. Crop yield and global food security will depend on how individual crop species respond to new and potentially more variable conditions. *B.rapa* is a laboratory crop model that has been successfully used to study the plant response to environmental change. phyB is emerging as a key regulator of carbon response and supply, metabolism and biomass production (Yang *et al.*, 2016; Arsovski *et al.*, 2018). In addition to a diminished high CO_2_ response we previously showed that *B.rapa phyB* mutants have reduced chlorophyll levels and photosynthetic rate (Arsovski *et al.*, 2018). In this work, we elicited a dark response in individual leaves and examined the transcriptomic response of wild-type and *phyB* leaves as they are darkened and were subsequently returned to light.

Wild-type *B.rapa* leaves darkened for three days have a significant upregulation of genes involved in autophagy, catabolism and leaf senescence while large groups of genes functioning in photosynthesis, metabolism and translation (Fig. 2A). This senescence response and redistribution of cell resources is typical of the dark response observed in various plant systems (Guo *et al.*, 2004; Brouwer *et al.*, 2012; Song *et al.*, 2014; Law *et al.*, 2018; Sobieszczuk-Nowicka *et al.*, 2018). Upon return to light upregulated and downregulated functional groups are essentially reversed. Genes acting mainly in response to light stimulus, photosynthesis, translation and metabolism were upregulated in leaves, while those with roles in catabolism, vesicle fusion and transport, and protein degradation were downregulated 24 hours following return to light (Fig. 3B). In many plants, leaf senescence is reversible in a limited time span after senescence initiation, leading to ‘regreening’ of the leaves. For example, a return to light following a 2 day dark period initiates a reconstitution of photosynthetic capability in *A. thaliana* (Parlitz *et al.*, 2011).

While there is a significant overlap in the transcriptomic response to dark and recovery between wild-type and *BrphyB* mutant leaves, there are important categories of genes that may explain some of phenotypes associated with the *BrphyB* mutant (Fig. 2,3). After three days of dark the orthologs of *A.thaliana* PHYTOCHROME INTERACTING FACTORS 4 and 5 are only upregulated in the *BrphyB* mutant suggesting a misregulation of the light response, as well as likely impacts on hormone homeostasis. In *A. thaliana* PIFs/EIN3/HY5-regulated genes in the dark were estimated to account for half of the light-directed transcriptome changes (Shi *et al.*, 2018). In recovery, *BrphyB* mutant leaves lack the increased transcription of genes involved in light harvesting and photosynthesis, including 125 chloroplast-targeted genes, and the mutants uniquely down-regulate genes involved in phototropism suggesting possible “crossed-wires” from mismatches between different photoreceptor responses (Fig. 3A). Together these finding may connect the observed reduction of chlorophyll and photosynthetic rate (Arsovski *et al.*, 2018) and fewer larger chloroplasts in *BrphyB* leaves compared to wild type.

Nitrogen is a critical resource that governs plant growth. The availability of nitrogen to the roots plays a particularly significant role in constraining plant growth and crop yield worldwide (Epstein and Bloom, 2005; Hirel *et al.*, 2011; Alvarez *et al.*, 2012). Nitrogen deficiency is one of the endogenous and environmental factors that regulates the onset of leaf senescence (Gregory, 1937; Mei and Thimann, 1984; Masclaux-Daubresse *et al.*, 2007; Koeslin-Findeklee *et al.*, 2015). In *B.rapa*, nitrogen availability limits growth increase in high CO_2_ (Arsovski *et al.*, 2018). In dark-induced leaf senescence nitrogen from senescing leaves is mobilized and transported to still growing vegetative tissues. Wild type *B.rapa* plants darkened for 3 days show a significant upregulation of genes involved in senescence, catabolism, and vesicle transport while downregulation of genes involved in protein synthesis indicating and active export of nitrogen resources (Supplementary File 1, 2). While *BrphyB* mutants largely share this response, there is evidence that this resource allocation is altered. After 3 days of dark phyB unique downregulated genes are significantly enriched for GO terms related to protein synthesis suggesting either a delayed or prolonged response compared to wild type. In recovery, *BrphyB* uniquely upregulated genes are significantly enriched for translation and peptide biosynthesis, while wild type unique genes are photosynthesis-related. *B.napus*, a close relative of *B.rapa* has poor nitrogen use efficiency (Masclaux-Daubresse *et al.*, 2007; Xu *et al.*, 2012). Only 50–60% of the applied nitrogen is recovered in the plants and at the time of harvest a relatively low 80% of the total plant nitrogen is localized in the seeds (Schjoerring *et al.*, 1995; Jensen *et al.*, 1997; Malagoli *et al.*, 2005; Rathke *et al.*, 2006). Here *BrphyB* mutants had a misregulation of genes orthologous to *A.thaliana* GLUTAMINE SYNTHASE 1;2, 1;3 and 1;4 that play a role in seed yield and size (Fig. 3C). While seeds at harvest were not significantly different from wild type in size and weight, seed yield is dramatically reduced in the mutant (Arsovski *et al.* 2018). A more detailed understanding of the phyB-regulated network holds the promise of improved plant growth models and identification of new targets for engineering more resource-efficient crops.

## Supporting information

Supplementary figure title page

## Supplemental Figures

Figure S1: qPCR validation.

Figure S2: Transcription factor motifs enriched in promoter regions.

Table S1: Gene expression comparison between BrphyB and wild type leaves in the Pre condition.

Table S2: Gene Ontology comparison of differentially expressed genes between Pre (P), Dark (D), and Recovery (R) in wild type and *BrphyB* leaves.

Table S3: A comparison of *A.thaliana* gene differentially expressed in whole darkened plants or individually darkened leaves in response to 3 days of darkness from Law et al.,2018 and *A.thaliana* orthologs of *B.rapa* genes differentially expressed in Dark vs. Pre.

Table S4: Gene Ontology analysis of *A.thaliana* genes differentially expressed in individually darkened leaves (IDL), or whole darkened plants (DP) from Law et al. 2018 and *A.thaliana* orthologs of *B.rapa* genes differentially expressed in Dark vs. Pre. Sheets show *A.thaliana* IDs, gene model names, MapMan bins and descriptions of common up or down regulated genes followed by GO annotations of those genes.

Table S5: A comparison of genes differentially expressed in dark and re-illumination from Parlitz et al. 2018 and *A.thaliana* orthologs of *B.rapa* genes in Dark vs. Pre and Recovery vs. Dark.

Table S6: Gene Ontology enrichment for DEGs common and unique to wild type and BrphyB in pre (P), dark (D), and recovery (R) conditions.

Table S7: Differentially expressed genes in pre, dark and recovery. Significance column denotes whether the significance is common to both wild type and *BrphyB* or unique to either.

## Acknowledgements

We thank Prof. Mark Stitt and Dr. Virginie Mengin for sharing their insights and expertise, as well as the other members of the PHYTOCAL consortium, Nemhauser and Imaizumi labs for feedback and discussion. We also thank the undergraduate researchers from Dr Arsovski’s Spring 2018 Special Field Topics class who carried out protocol optimization and preliminary measurements of phenotypes presented here: Ericka Budinich, Jonas Hill, Sean Hoeger, Katrin Hosseini, Nikhil Kaza, Winnie Kwong, Kellen Larsen, Andrew Lui, Rohan Menon, Claudia Moroney, Anita Nguyen, Arthur Sargent, Emma Stevens, and Nanami Tsumura. This work was supported by the National Science Foundation participation in the ERA-CAPS program (IOS-1539834).

