## Supplementary figure title page for "*BrphyB* is critical for rapid recovery to darkness in mature *Brassica rapa* leaves"

**Supplemental Figures**

Figure S1: qPCR expression mirrors RNAseq for select genes. Left, RNAseq expression of the 3 biological replicates. Right, qPCR of the same genes on left using 3 different biological replicates. Error bars are SE.

Figure S2: (A) Logo plots of motifs significantly enriched in recovery vs. dark, clustered by sequence similarity. (B) percent of genes differentially regulated in recovery. (C) Frequency of TELO-box motif occurrence in recovery (R) versus dark (D) differentially expressed genes. (D) Frequency of Evening Element motif occurrence in recovery (R) versus dark (D) differentially expressed genes.

Table S3: A comparison of *A.thaliana* gene differentially expressed in whole darkened plants or individually darkened leaves in response to 3 days of darkness from Law et al.,2018 and *A.thaliana* orthologs of *B.rapa* genes differentially expressed in Dark vs. Pre.
