## Supplemental Figure 1 for "*BrphyB* is critical for rapid recovery to darkness in mature *Brassica rapa* leaves"

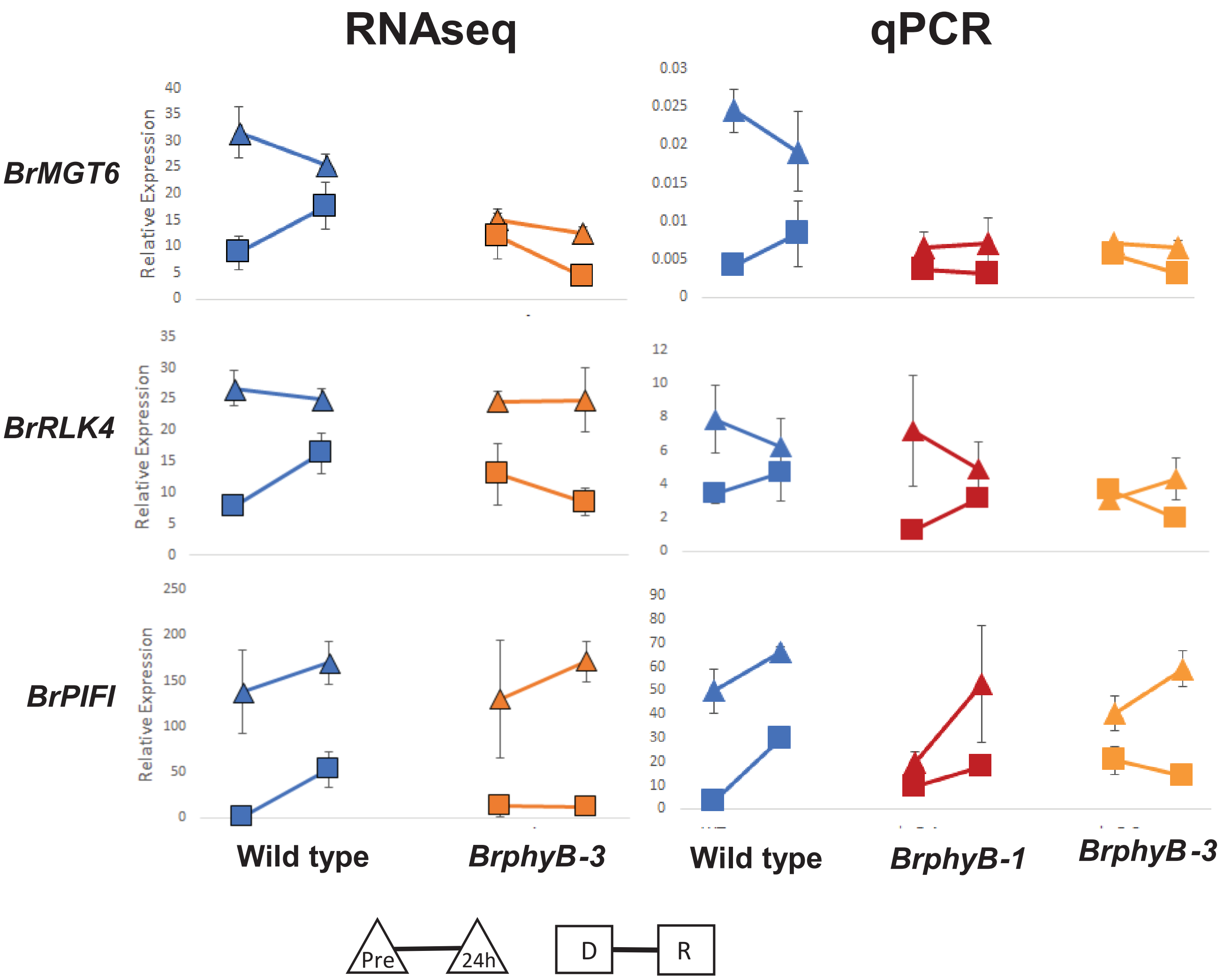


**Supplementary Figure 1:** qPCR expression mirrors RNAseq for select genes. Left, RNAseq expression of the 3 biological replicates. Right, qPCR of the same genes on left using 3 different biological replicates. Error bars are SE .
