## Supplemental Figure 2 for "*BrphyB* is critical for rapid recovery to darkness in mature *Brassica rapa* leaves"

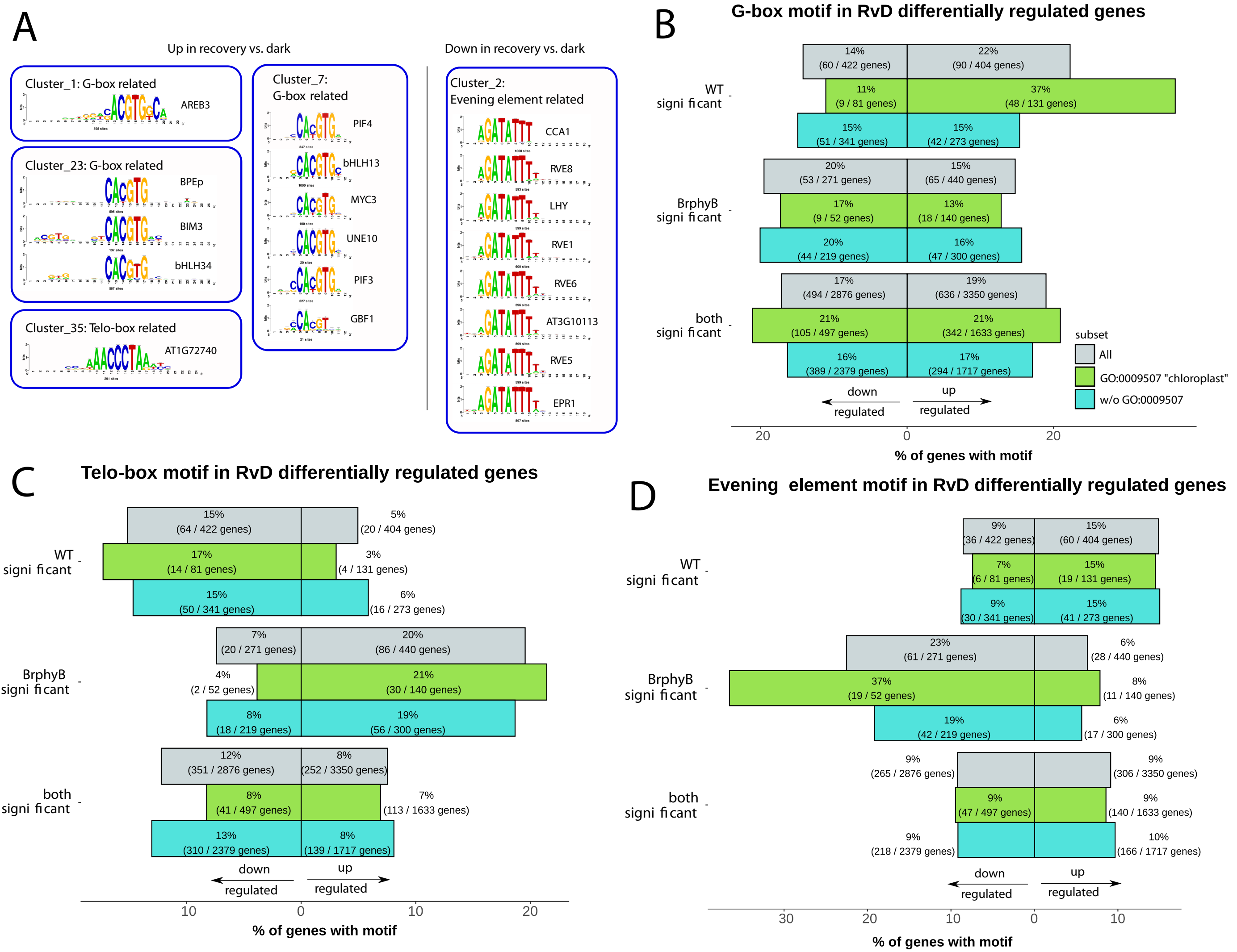


**Supplemental Figure S2:** (A) Logo plots of motifs significantly enriched in recovery vs. dark, clustered by sequence similarity. (B) percent of genes differentially regulated in recovery. (C) Frequency of TELO-box motif occurrence in recovery (R) versus dark (D) differentially expressed genes. (D) Frequency of Evening Element motif occurrence in recovery (R) versus dark (D) differentially expressed genes.
