## Supplemental Table 3 for "*BrphyB* is critical for rapid recovery to darkness in mature *Brassica rapa* leaves"

| | Whole plant | Individual leaves | Both |
|---|---|---|---|
| Law et al.2018UP DP/IDL only | 592 | 1057 | 2042 |
| Law et al.2018DWN DP/IDL only | 661 | 779 | 3040 |
| Law et al.2018TOT DP/IDL only | 1253 | 1836 | 5082 |
| Common UP | 147 | 385 | 951 |
| Common DWN | 168 | 263 | 1626 |
| Common Total | 315 | 648 | 2577 |
| % Common UP | 24.83% | 36.42% | 46.57% |
| % Common DWN | 25.42% | 33.76% | 53.49% |
| % Common Total | 25.14% | 35.29% | 50.71% |
| % random chance UP | 21.91% | 21.91% | 21.91% |
| % random chance DWN | 25.23% | 25.23% | 25.23% |
| % random chance both | 23.66% | 23.32% | 23.89% |

**Supplementary Table 3:** A comparison of *A.thaliana* gene differentially expressed in whole darkened plants or individually darkened leaves in response to 3 days of darkness from Law et al.,2018 and *A.thaliana* orthologs of *B.rapa* genes differentially expressed in Dark vs. Pre.
